# G-CSF and IL-6 drive myeloid dysregulation during severe viral infection

**DOI:** 10.1101/2024.09.12.612764

**Authors:** Kimberly Kajihara, Donghong Yan, Gretchen L. Seim, Hannah Little-Hooy, Jing Kang, Cynthia Chen, Marco De Simone, Tim Delemarre, Spyros Darmanis, Haridha Shivram, Rebecca N. Bauer, Carrie M. Rosenberger, Sharookh B. Kapadia, Min Xu, Miguel Reyes

## Abstract

Dysregulated myeloid states are associated with disease severity in both sepsis and COVID-19. However, their relevance in non-COVID-19 viral infection, the factors driving their induction, and their role in tissue injury remain poorly understood. We performed a meta-analysis of 1,622,180 myeloid cells from 890 COVID-19 or sepsis patients and controls across 19 published blood scRNA-seq datasets, which revealed severity-associated gene programs in both neutrophils and monocytes pointing to emergency myelopoiesis (EM). Using published bulk transcriptional data from 562 individuals with non-COVID-19 viral disease, we show that these signatures are similarly upregulated during severe influenza and RSV infection. Analysis of transcriptional and proteomic responses in tocilizumab-treated COVID-19 patients show that IL-6 signaling blockade results in a partial reduction of EM signatures and a compensatory increase in the growth factor G-CSF. Using a cellular model of human myelopoiesis, we show that both IL-6 and G-CSF stimulate the production of myeloid cells that express EM signatures *in vitro*. Using a mouse model of severe influenza infection, we demonstrate the effect of IL-6 and G-CSF signaling blockade on EM-associated myeloid cells, and highlight the opposing effects of EM-induced neutrophils and monocytes on tissue injury. Our study demonstrates the link between systemic cytokines and myeloid dysregulation during severe infection in humans, and highlights the cooperative role of IL-6 and G-CSF signaling in driving infection-induced myelopoiesis.

## INTRODUCTION

Respiratory viruses have caused four major epidemics, including the 2009 Influenza A/H1N1 and COVID-19 pandemics, in the last two decades (*1*). These pandemics highlight the importance of accelerating therapeutic development for viral respiratory diseases to mitigate future outbreaks. Virus-directed therapies are often the first-line strategy against respiratory infections; however, existing antivirals have a narrow spectrum of activity (*2*) and are often ineffective in patients with severe disease (*3–5*). In contrast, host-directed therapies can be pathogen-agnostic, and have recently demonstrated efficacy in patients with severe COVID-19 (*6*). Understanding the pathogenesis of severe viral infection and the specific role of the immune response are critical to developing host-directed therapies (*7*).

Several studies have reported profound transcriptional and immunophenotypic alterations, particularly in myeloid cells, in patients with severe viral disease (*8–13*). These studies highlight the expansion of immature monocytes, neutrophils, and myeloid progenitors in circulation, commonly referred to as emergency myelopoiesis (EM) (*14*). We previously described the expansion of MS1 monocytes (IL1R2+HLA-DR^low^) in both bacterial sepsis and severe COVID-19 (*15*, *16*). These cells can be induced *in vitro* by treatment of human hematopoietic stem and progenitor cells (HSPCs) with plasma from patients with sepsis or severe COVID-19 (*15*), indicating a role for EM in its induction. Recently, EM has also been proposed as a driver of severity-associated *IL1R2*^+^ immature neutrophils in sepsis and COVID-19 (*17–19*). Despite these reports, the relevance of these cell states in non-COVID-19 viral infection and their impact on tissue injury remain unclear. Here, we show that transcriptional signatures of EM are associated with disease severity in influenza and RSV infection, in addition to bacterial sepsis and COVID-19. We identified cytokines that drive these signatures in humans, and characterized the role of EM-associated monocytes and neutrophils on tissue injury *in vivo*.

## RESULTS

### Large-scale meta-analysis identifies shared monocyte and neutrophil gene programs in sepsis and severe COVID-19

We previously performed a metaanalysis of blood monocytes from 5 scRNA-seq datasets to demonstrate the expansion of MS1 monocytes in both sepsis and severe COVD-19 (*15*). To validate these findings and identify shared transcriptional programs across myeloid cells, we expanded this analysis to 18 datasets (*9*, *10*, *15–30*), including 6 studies with transcriptional data from blood neutrophils (**Figure 1a, Methods, Supplementary Table 1**). Our expanded meta-analysis of monocytes identified gene programs shared across datasets, including the MS1 and MHC-II gene programs (**Supplementary Figure 1**). Consistent with our previous findings, we found that, across all datasets, usage of the MS1 and MHC-II gene programs are increased and decreased, respectively, in sepsis patients compared to healthy controls, and in severe compared to mild COVID-19 (**Figure 1b-c, Supplementary Figure 2**, **Supplementary Table 2,4**).

**Figure 1.**
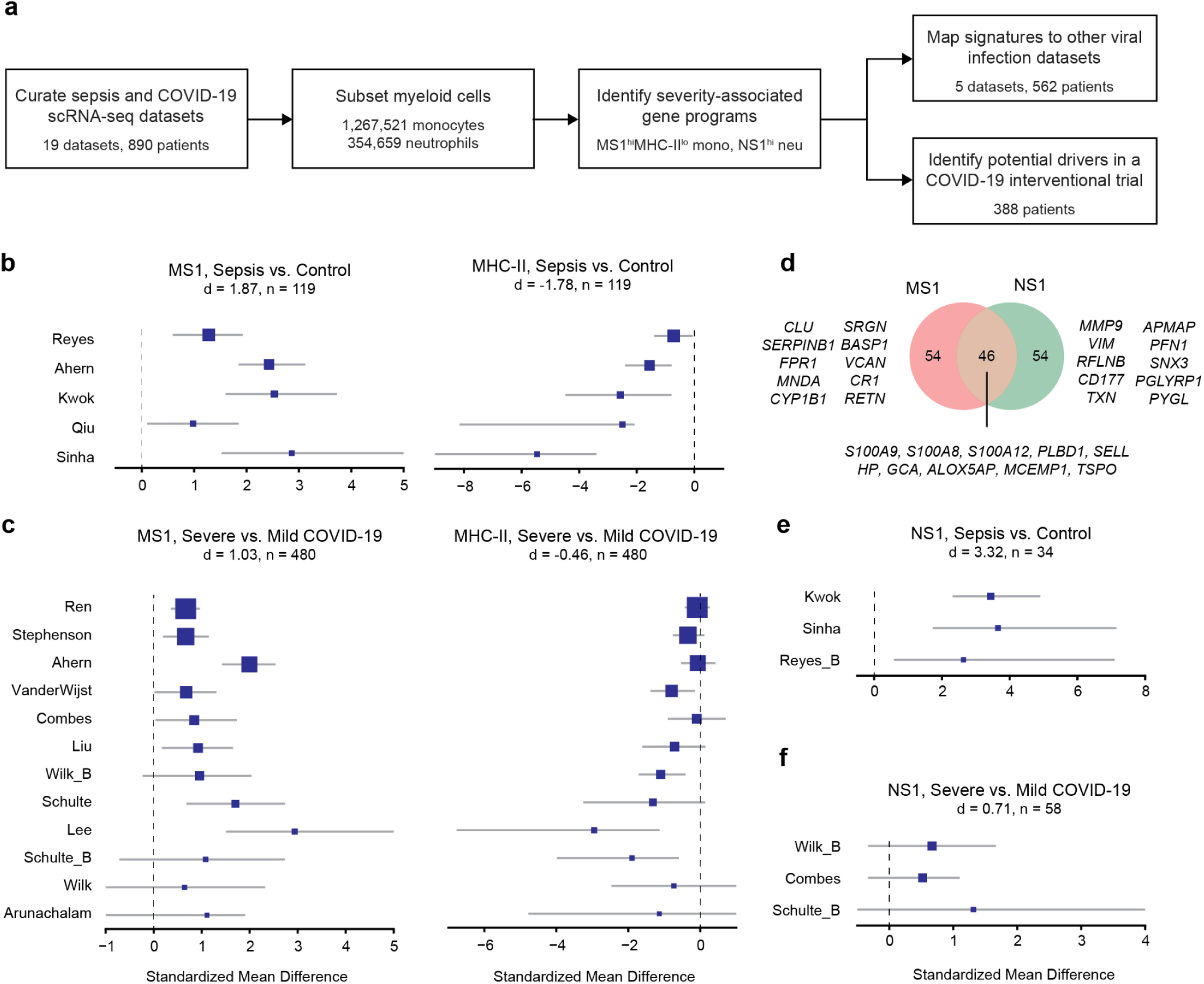
Emergency myelopoiesis gene programs are associated with sepsis and COVID-19 severity. (a) Analysis scheme for 18 scRNA-seq datasets and 5 bulk transcriptomics datasets from cohorts of patients with bacterial sepsis, COVID-19, or non-COVID-19 viral infection. (b-c, e-f) Forest plots indicate the effect sizes (log2 standardized mean difference) across datasets for each indicated gene program. Blue boxes indicate the effect size in an individual study, with whiskers extending to the 95% confidence interval. Size of each box is proportional to the relative sample size of the study. Detailed descriptions of the patient cohorts and numbers of cells and patients for each dataset are described in the corresponding publications (*9*, *10*, *15–30*) and listed in **Supplementary Table 1**. (d) Venn diagram showing the overlap between the top 100 genes in the MS1 and NS1 programs. The top ten genes with highest loadings in each region are indicated.

We performed a metaanalysis of blood neutrophils (**Supplementary Figure 3a-d**), and identified a neutrophil program, which we named NS1, that shares several genes with the MS1 monocyte program (**Figure 1d, Supplementary Table 3**). NS1 is similarly upregulated in sepsis and severe COVID-19, though with a smaller effect size when comparing mild vs. severe COVID-19 (**Figure 1e-f, Supplementary Figure 3c-d, Supplementary Table 4**). Usage of the MS1 and NS1 gene programs across patients correlated significantly in 3 out of 5 datasets with both cell types, suggesting that these gene programs may be driven by similar pathways (**Supplementary Figure 3e**). Of note, the top genes of the NS1 program also point to EM, as the expression of granule genes in blood neutrophils are indicative of premature release of developing neutrophils (*31*). Our metaanalysis shows that EM gene programs can be detected in both monocytes and neutrophils, and their increased expression in both cell types is associated with sepsis and COVID-19 severity.

### EM signatures are associated with disease severity in non-COVID-19 viral infection

Given the increased expression of EM-associated gene programs in both sepsis and severe COVID-19, we hypothesized that similar patterns will be observed in non-COVID-19 respiratory infections. We used the top genes associated with each consensus gene program from our meta-analysis (MS1, MHC-II, and NS1) to estimate the expansion of EM-associated cell states in bulk transcriptional datasets (**Methods, Supplementary Table 5**). Interestingly, we found that EM scores significantly increase with disease severity in 4 out of 5 datasets (**Figure 2a-b**), with a summary effect size of 1.58 across all datasets (**Figure 2c**, **Supplementary Table 6**). Consistent with this result, usage of MS1 and MHC-II are increased and decreased, respectively, in two scRNA-seq datasets that included cohorts of patients with severe influenza (**Supplementary Figure 1c-d, 2a-b**). These results suggest that the severity-associated increase in EM signatures is a conserved response across viral infections, regardless of etiology.

**Figure 2.**
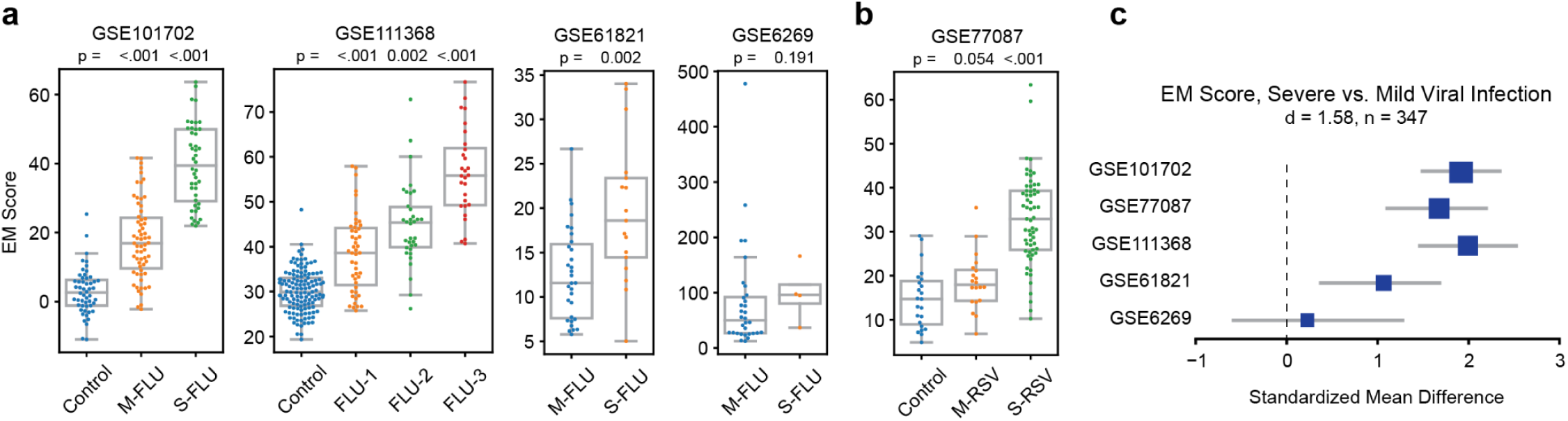
Emergency myelopoiesis increases with severity in non-COVID-19 viral infection. (a-b) EM scores in blood transcriptional datasets of influenza (a) and RSV (b) patients. Boxes show the median and interquartile range (IQR) for each patient cohort, with whiskers extending to 1.5 IQR in either direction from the top or bottom quartile. Accession numbers for each dataset are indicated above. p values shown are calculated by comparing each disease state with Control (GSE101702 and GSE111368) or M-FLU group (GSE61821 and GSE6269) using a two-tailed Wilcoxon rank-sum test. Detailed descriptions of the patient cohorts and numbers of patients for each dataset are described in the corresponding publications (*11*, *12*, *49–51*) and listed in **Supplementary Table 5**. (c) Forest plots indicate the effect sizes (log2 standardized mean difference) of the EM score between mild vs. severe patients across datasets. Accession numbers for each dataset are indicated on the left. Blue boxes indicate the effect size in an individual study, with whiskers extending to the 95% confidence interval. Size of each box is proportional to the relative sample size of the study. M-FLU, mild influenza; S-FLU, severe influenza; FLU-1, influenza with no supplemental oxygen required; FLU-2, influenza with oxygen by mask required, FLU-3, influenza with mechanical ventilation. M-RSV, mild respiratory syncytial virus infection; S-FLU, severe respiratory syncytial virus infection.

### EM signatures are partially reduced by IL6 signaling blockade

We previously demonstrated that the induction of MS1 monocytes in HSPCs by sepsis plasma is partially mediated by IL-6 (*15*). To confirm the role of IL-6 in MS1 induction in COVID-19 patients, we analyzed published bulk transcriptional data from COVACTA: a phase 3 randomized placebo-controlled study of tocilizumab, an anti-IL-6R antibody, in hospitalized COVID-19 patients (*8*, *32*). Consistent with our findings, we found that EM scores are correlated with baseline ordinal severity scores (**Supplementary Figure 4a**). In addition, higher baseline EM scores were significantly associated with mortality and mechanical ventilation on day 28 in placebo-treated patients (PBO; **Figure 3a**). Importantly, we found that tocilizumab-treated patients (TCZ) have a greater reduction in EM scores compared with the PBO group on days 3 and 7 after treatment (**Figure 3b**), confirming the role of IL-6 signaling in inducing EM in COVID-19 patients. Of note, patients who are still mechanically ventilated (MV) at day 28 had significantly higher EM scores, independent of treatment, indicating that IL-6 signaling blockade is not sufficient to completely inhibit EM in patients with critical COVID-19.

**Figure 3.**
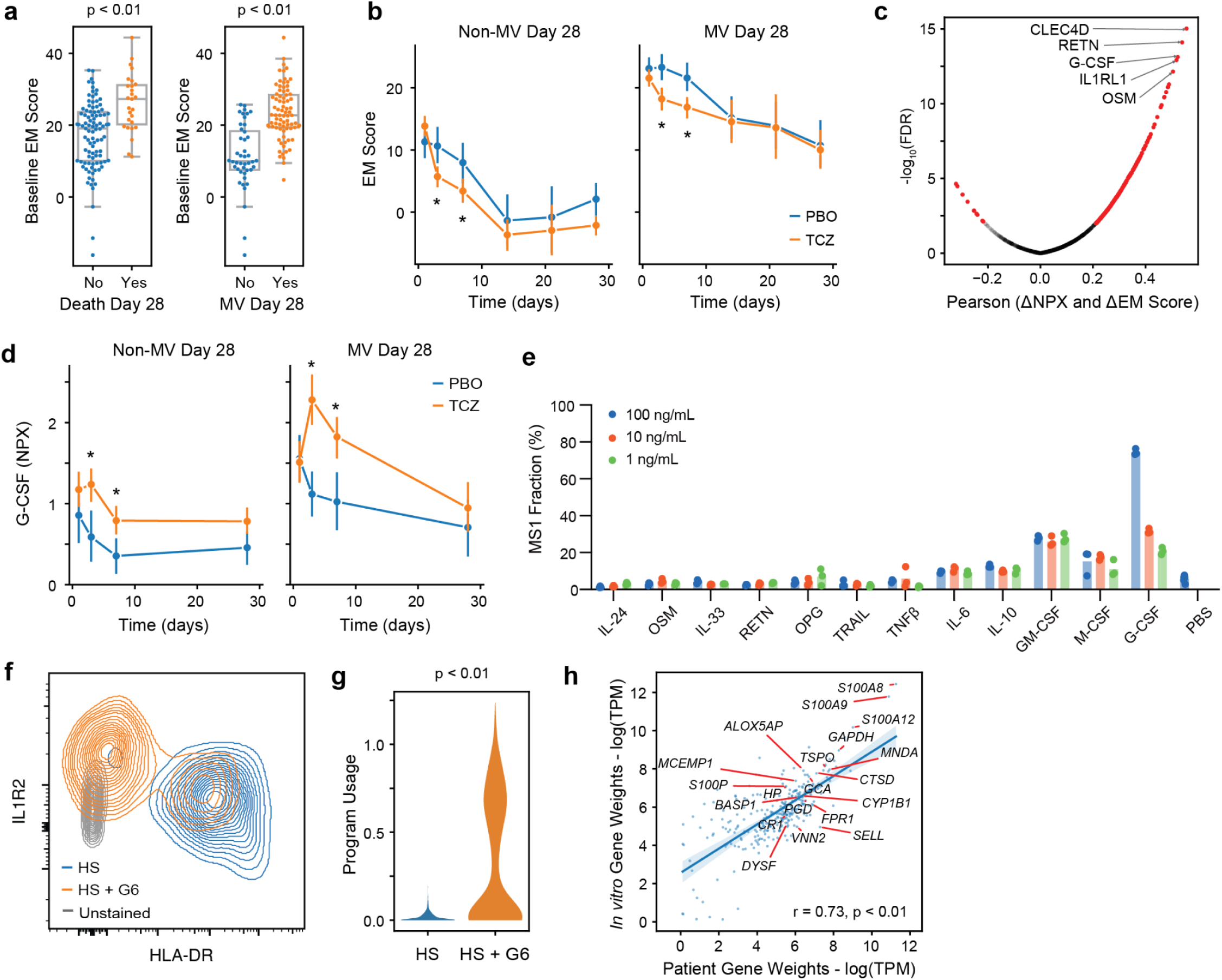
IL-6 and G-CSF drive emergency myelopoiesis and production of MS1 monocytes. (a) EM scores in patients from the PBO arm of COVACTA grouped by mortality (left) or mechanical ventilation status (right) on day 28. Boxes show the median and interquartile range (IQR) for each patient cohort, with whiskers extending to 1.5 IQR in either direction from the top or bottom quartile. p values shown are calculated for each comparison using a two-tailed Wilcoxon rank-sum test. (b) EM scores over time in patients from the PBO or TCZ arm of COVACTA, grouped by mechanical ventilation status. EM scores are averaged across all patients for each group and timepoint. Error bars indicate the 95% confidence interval of the mean. Asterisks indicate a false discovery rate (FDR) < 0.05, computed by comparing the PBO and TCZ group at each timepoint (two-tailed Wilcoxon rank sum test, corrected for multiple testing). (c) Volcano plot showing the Pearson correlation between changes in protein levels and changes in EM score from baseline to day 3 in the TCZ group. Significance of each correlation was calculated with a two-sided permutation test. Proteins with FDR of <0.01 are highlighted in red. (d) G-CSF levels over time in patients from the PBO or TCZ arm of COVACTA, grouped by mechanical ventilation status. G-CSF levels are averaged across all patients for each group and time point. Error bars indicate the 95% confidence interval of the mean. Asterisks indicate a false discovery rate (FDR) < 0.05, computed by comparing the PBO and TCZ group at each timepoint (two-tailed Wilcoxon rank sum test, corrected for multiple testing). (e) Fraction of MS1 (IL1R2+HLA-DR^low^) among monocytes in HSPCs incubated with each cytokine at the indicated concentration. (f) Flow cytometry density plot of CD14+ monocytes from HSPCs treated with healthy serum (HS) or healthy serum with 10 ng/mL IL-6 and G-CSF (HS+G6). (g) Violin plot of MS1 gene program usage in monocytes differentiated from HSPCs treated with HS or HS+G6. (h) Gene weight correlation between the MS1 program detected in patients (x-axis) and the MS1 program detected in differentiated HSPCs (y-axis). Significance of the Pearson correlations (r) was calculated with a permutation test. The top 20 genes with the highest z-score loadings in the patient MS1 program are labeled. NPX, normalized protein expression.

To identify other potential EM drivers besides IL-6, we analyzed correlations between blood protein levels and leukocyte gene expression in TCZ-treated patients. We identified several proteins whose increase in expression from baseline to day 3 are significantly correlated with increases in EM scores (**Figure 3c, Supplementary Table 7**). Among the top proteins was G-CSF, a cytokine that plays a central role in granulocyte production (*33*, *34*). Of note, G-CSF levels significantly increase with TCZ treatment in COVACTA as we reported previously (*32*), and changes in both G-CSF and EM scores are higher in patients who are mechanically ventilated at day 28 (**Figure 3d, Supplementary Figure 4b**). In addition, while both changes in IL-6 and G-CSF correlate with EM in the PBO group, G-CSF has a stronger correlation with EM in the TCZ group (**Supplementary Figure 4c**). These data suggest that G-CSF is a potential driver of EM in severe COVID-19, and that IL-6 signaling blockade may result in a compensatory increase in G-CSF that maintains elevated EM, particularly in critical COVID-19 patients.

### G-CSF and IL-6 induce neutrophil and MS1 monocyte production

To test candidate EM drivers, we incubated human HSPCs with cytokines that correlated with EM in our analysis of COVACTA, and measured their effect on total monocyte, MS1, and neutrophil production using flow cytometry. We tested these proteins along with IL-6, IL-10, GM-CSF, and M-CSF, which we previously showed induces MS1 production in combination (*15*). Among the cytokines we tested, G-CSF produced the highest number of neutrophils and monocytes, and specifically induced monocytes with the MS1 immunophenotype (IL1R2+HLA-DR^low^; **Figure 3e, Supplementary Figure 5a**). Interestingly, while higher concentrations of G-CSF and IL-6 induced greater MS1 production in combination, IL-6 slightly reduced neutrophil production in the presence of G-CSF (**Figure 3f, Supplementary Figure 5b**).

We performed scRNA-seq on HSPCs differentiated with IL-6 and G-CSF to confirm the induction of MS1 monocytes by gene expression (**Supplementary Figure 6a**). We found increased and decreased expression of MS1 and MHC-II genes, respectively, in CD14+ cells generated from HSPCs with IL-6 and G-CSF compared with healthy serum alone (**Supplementary Figure 6b-c**). Unbiased analysis of the *in vitro* scRNA-seq data also detected gene programs similar to MS1 and MHC-II in patients, and their usage patterns are consistent with those observed in sepsis and COVID-19 (**Figure 3g-h, Supplementary Figure 6d-f).** Altogether, these data suggest that both G-CSF and IL-6 could induce EM signatures observed in patients, and that their combination is sufficient to produce MS1 monocytes *in vitro*.

### EM signatures are recapitulated by influenza infection in mice

To understand the effects of emergency myelopoiesis on tissue injury and disease severity *in vivo*, we utilized a mouse model of influenza A/PuertoRico/8/34 H1N1 (PR8) infection. Given the association of EM signatures with influenza severity in humans (**Figure 2a**), we first evaluated the relationship between EM signatures and infection dose (**Figure 4a**). While we did not find dose-dependent differences in the absolute number of monocytes in the blood and BALF, we found a specific increase in IL1R2+ monocytes (**Figure 4b-d, Supplementary Figure 7a**). IL1R2 has also been proposed as a specific marker of immature neutrophils in sepsis and severe COVID-19 patients (*17*, *19*). Consistent with this, we found a significant increase in IL1R2 expression among neutrophils with higher infection dose (**Figure 4e-f, Supplementary Figure 7b**). These data suggest that monocytes and neutrophils at higher infection doses are similar to those found in patients with severe infection.

**Figure 4.**
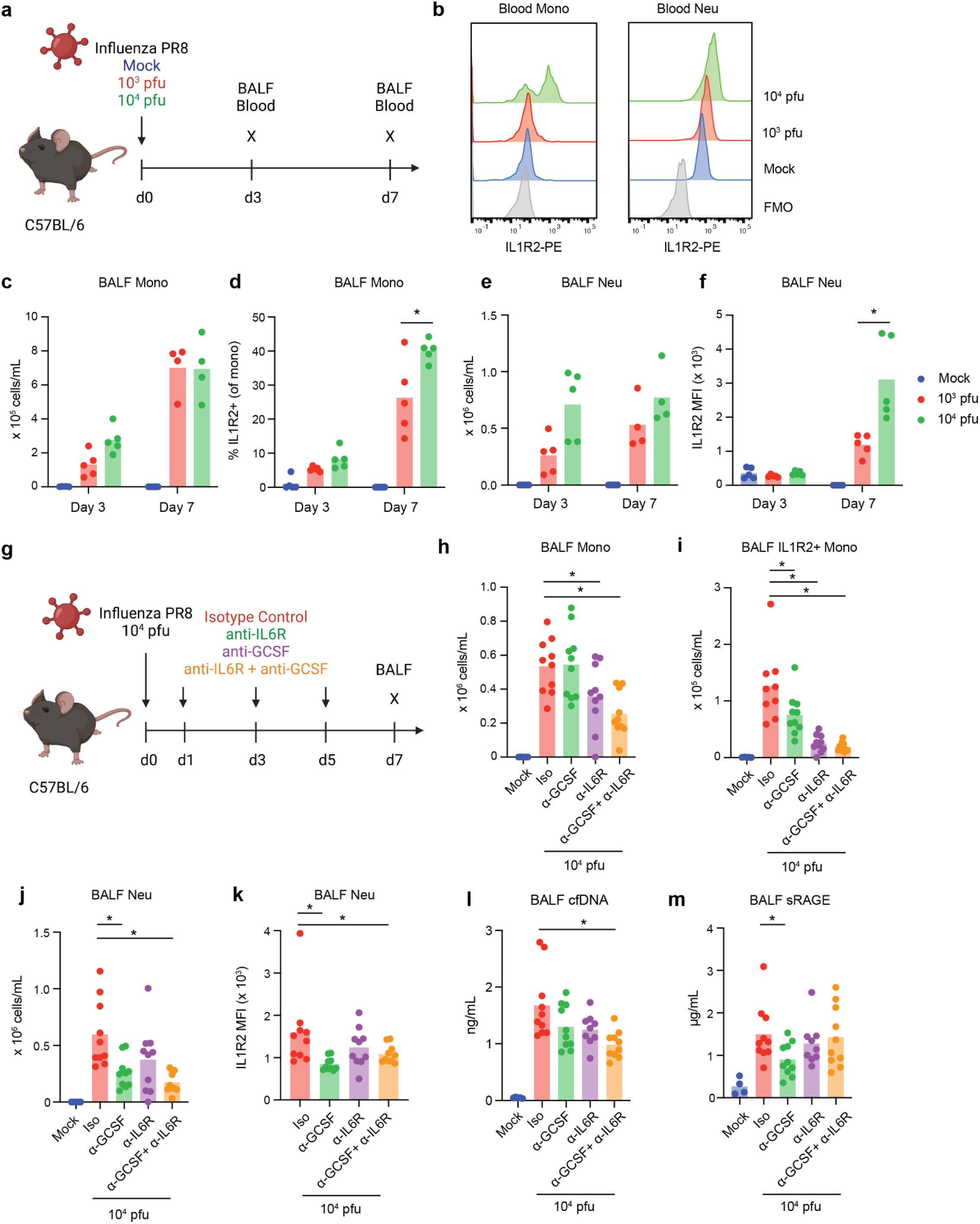
IL-6 and G-CSF drive emergency myelopoiesis during severe influenza infection *in vivo*. (a) Schematic of the influenza dose titration experiment. (b) IL1R2 surface expression at day 7 post-infection in monocytes (left) and neutrophils (right) in the blood of mice infected with the indicated PR8 titer. (c) Number and (d) fraction of IL1R2+ blood monocytes in mice infected with PR8 at the indicated time point. (e) Number and (f) IL1R2 mean fluorescence intensity (MFI) of blood neutrophils in mice infected with PR8 at the indicated time point. (g) Schematic of the IL6 and G-CSF blockade experiment. (h) Number of total monocytes, (i) IL1R2+ monocytes, (j) neutrophils, and (k) neutrophil IL1R2 mean fluorescence intensity (MFI) at day 7 post-infection in the BALF of mice infected with 10^4^ PR8 and treated with the indicated antibodies. (l) cfDNA and (m) sRAGE levels at day 7 post-infection in the BALF of mice infected with 10^4^ PR8 and treated with the indicated antibodies. Asterisks indicate a significant difference (p < 0.05, unpaired t-test). BALF, bronchoalveolar lavage fluid; Iso, isotype control.

To confirm the presence of EM signatures *in vivo* by gene expression, we performed scRNA-seq on blood and BALF immune cells from PR8-infected mice (**Supplementary Figure 8a-b**). Prior to analysis, we converted mouse to human genes in the scRNA-seq data to enable the quantification of our human signatures *in vivo* (**Methods**). Similar to the patterns observed in humans, we found increasing expression of the MS1 and NS1 genes in mouse monocytes and neutrophils, respectively, with higher infection dose (**Supplementary Figure 8c**). In addition, unbiased cNMF analysis revealed gene programs similar to MS1 and NS1, and their usages increased with higher infection dose (**Supplementary Figure 8d-g**). Similar to patients with severe COVID-19, systemic levels of G-CSF and IL-6 are elevated with PR8 infection, and their levels correlate across individual mice (**Supplementary Figure 7c-d**). Altogether, these data suggest that EM signatures and their severity-associated increases in humans can be modeled by high-dose PR8 infection in mice.

### Combined IL-6 and G-CSF signaling blockade inhibits EM in vivo

Given the associations of IL-6 and G-CSF with EM signatures in patients and their effect in inducing EM *in vitro*, we tested the effects of blocking these cytokines or their receptors during high-dose PR8 infection *in vivo* (**Figure 4g**). Treatment with α-G-CSF or α-IL-6R significantly reduced either total neutrophil or monocyte numbers, respectively, while combined treatment resulted in a significant decrease in both cell types (**Figure 4h,i**). Both α-G-CSF and α-IL-6R significantly reduced the number of IL1R2+ monocytes, and their combination resulted in a greater numerical decrease (**Figure 4j**). In addition, α-G-CSF and its combination with α-IL-6R reduced the expression of IL1R2 in BALF neutrophils (**Figure 4k**). These findings confirm the role of IL-6 and G-CSF signaling in inducing both EM-associated monocytes and neutrophils during severe influenza infection *in vivo*.

To assess the consequence of targeting EM on tissue injury *in vivo*, we measured biomarkers of broad tissue injury (cell-free DNA) (*35*, *36*) and alveolar injury (sRAGE) (*37*, *38*) in the BALF. Individual treatment with α-G-CSF or α-IL-6R resulted in modest reduction of cfDNA levels, however, combined treatment significantly reduced cfDNA levels (**Figure 4l**). Interestingly, we found that treatment with α-G-CSF, but not α-IL6-R or their combination, significantly decreased sRAGE levels in the BALF (**Figure 4m**). These mixed results suggest that inhibiting EM may have a limited effect in preventing tissue injury, perhaps due to differing roles of EM-associated monocytes and neutrophils.

### Cell type-specific effects of EM in vivo

To understand the role of EM-induced monocytes *in vivo*, we performed high-dose PR8 infection in Ccr2-/- mice (**Figure 5a**). Ccr2 deletion resulted in significant decreases in numbers of both total monocytes and IL1R2+ monocytes, but not neutrophils (**Figure 5b-d**). Strikingly, we found that PR8 infection in Ccr2-/- mice resulted in increased levels of both tissue injury markers, cfDNA and sRAGE (**Figure 5e-f**). These increases are not associated with defects in pathogen control, as viral titers are similar between WT and Ccr2-/- mice (**Figure 5g**).

**Figure 5.**
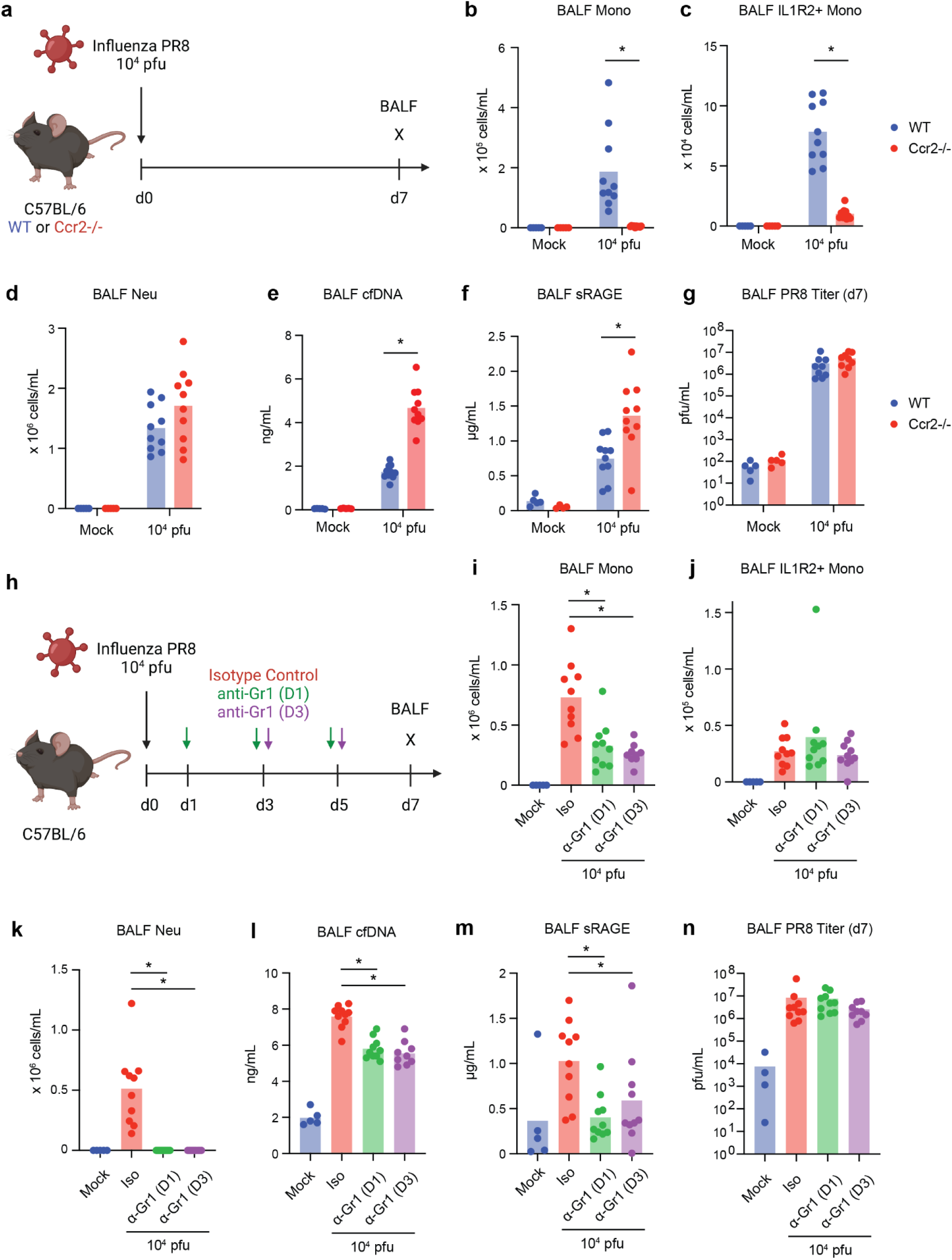
Opposing effects of monocytes and neutrophils on tissue injury during severe influenza infection *in vivo*. (a) Schematic of the Ccr2-/- influenza experiment. (b) Number of total monocytes, (c) IL1R2+ monocytes and (d) neutrophils at day 7 post-infection in the BALF of wild type (blue) or Ccr2-/- (red) mice infected with 10^4^ pfu PR8. (e) cfDNA, (f) sRAGE, and (g) PR8 levels at day 7 post-infection in the BALF of wild type (blue) or Ccr2-/- (red) mice infected with 10^4^ pfu PR8. (h) Schematic of the neutrophil depletion experiment. (i) Number of total monocytes, (j) IL1R2+ monocytes and (k) neutrophils at day 7 post-infection in the BALF of mice infected with 10^4^ pfu and treated with α-Gr1. (l) cfDNA, (m) sRAGE, and (n) PR8 levels at day 7 post-infection in the BALF of mice infected with 10^4^ pfu and treated with α-Gr1. Asterisks indicate a significant difference (p < 0.05, unpaired t-test). BALF, bronchoalveolar lavage fluid; Iso, isotype control.

Conversely, we used antibody-mediated depletion to understand the role of neutrophils during EM (**Figure 5h**). α-Gr1 treatment initiated on days 1 or 3 post-infection resulted in near-complete elimination of neutrophils on day 7 (**Figure 5k**). anti-Gr1 also resulted in partial reduction of monocyte numbers; however, the number of IL1R2+ monocytes were unaffected (**Figure 5i-j**). We found significant reductions in cfDNA and sRAGE levels in anti-Gr1 treated mice without concomitant changes in viral titer (**Figure 5l-n**).

Together, these results show that neutrophils are the primary mediators of tissue damage, while monocytes play a beneficial role, during EM. Importantly, these findings suggest that completely inhibiting EM may not be efficacious, as it may limit not only tissue injury caused by neutrophils, but also the beneficial effect of IL1R2+ monocytes.

## DISCUSSION

Our study demonstrates that EM-associated cell states are expanded across severe bacterial and viral infections in humans. By performing a meta-analysis of published single cell and bulk transcriptional datasets, we show that gene programs associated with EM are upregulated in sepsis patients and correlated with severity in patients with SARS-CoV-2, RSV, or influenza infection. These findings are consistent with a recent report identifying conserved severity-associated gene modules, including myelopoiesis, across multiple viruses (*39*). Quantification of EM gene programs in patients with non-COVID-19 viral infection was limited to bulk transcriptional datasets in our study, and additional scRNA-seq data is needed to specifically confirm the up-regulation of EM gene programs in neutrophils and monocytes from influenza and RSV-infected patients.

Using data from a previous clinical trial and *in vitro* experiments, we show that EM-associated cell states can be induced by G-CSF and IL-6. In addition, we show that combined targeting of G-CSF and IL-6 reduces EM-associated monocytes and neutrophils in a mouse model of severe influenza infection. Both cytokines have been proposed as prognostic biomarkers for influenza-associated pneumonia in humans (*40*). While G-CSF levels did not correlate with baseline severity or mortality in COVACTA, G-CSF levels at baseline were significantly associated with mechanical ventilation at day 28 (**Supplementary Figure 4d**). Our analysis also highlights the compensatory increase in G-CSF in patients treated with TCZ and its correlation with EM scores. The mechanism driving this response in humans, including the cell types that respond to IL-6R blockade and lead to increased G-CSF secretion, remains unknown. In addition, the specific bone marrow precursors that are mobilized by G-CSF and IL-6 and produce IL1R2+ monocytes or IL1R2^hi^ neutrophils remain to be identified.

Using Ccr2-deficient mice and antibody-mediated neutrophil depletion, we show the opposing effects of monocytes and neutrophils on severe infection-induced tissue injury, independent of their role in pathogen control. Greater neutrophil infiltration and activation in the airways are associated with disease severity in COVID-19 (*41*, *42*), and pre-clinical studies have demonstrated the role of neutrophil extracellular traps in driving tissue injury (*43*, *44*). Given the tissue-damaging effects of neutrophils, the production of immunosuppressive IL1R2+ monocytes perhaps acts as a negative feedback mechanism to limit neutrophil-mediated tissue injury. Consistent with this hypothesis, it has been reported that G-CSF and IL-6 also induce monocytic myeloid-derived suppressor cells (M-MDSCs) in the context of cancer and allotransplantation (*45*, *46*). While our results demonstrate the opposing roles of total monocytes and neutrophils during severe infection *in vivo*, the effect of specifically depleting IL1R2+ monocytes or IL1R2^hi^ neutrophils needs to be determined. Nevertheless, our findings highlight the importance of cell-specific targeting to modulate the host response during infection, as myeloid cells may play both detrimental and protective roles.

Overall, our study reveals the drivers of dysregulated myeloid cells during severe infection and their contribution to tissue injury. Our meta-analysis across various infectious diseases suggests that these responses are conserved across multiple pathogens, and that findings from our study may be leveraged to guide diagnostic and therapeutic development for future outbreaks.

## METHODS

### Meta-analysis of published scRNA-seq and bulk transcriptional datasets

Filtered gene expression matrices, including their patient and cohort annotations, were obtained from each publication (**Supplementary Table 1**) and analyzed using scanpy v1.9.1 and cNMF v1.4.1. We first subsetted monocytes and/or neutrophils from each dataset by using annotations provided by each study. If annotations were unavailable, we performed Leiden clustering and identified monocyte and neutrophil clusters by expression of *LYZ, CD14, CSF3R,* and *S100A8*. After subsetting, we performed cNMF on each individual gene expression matrix with the following parameters: k = 15, n-iter = 10, numgenes = 5000 and local-density-threshold = 0.2. To exclude gene programs from contaminating cell types or those which are lowly expressed, we employed an additional filter and removed programs whose mean usage across all cells is < 0.01.

Once cNMF results were obtained from each individual dataset, we identified common gene programs by concatenating and clustering the gene spectra matrices from all datasets. We used hierarchical clustering (scipy v1.10.0) on the concatenated gene spectra matrix with the following parameters: method = ‘average’, metric = ‘correlation’. To identify the top genes in each program cluster, we averaged the gene spectra scores across the programs and ranked the genes by these values. We identified the MS1 and NS1 gene programs by the presence of the genes *S100A8, S100A9, S100A12,* and *ALOX5AP* among the top 10. Similarly, we identified the MHC-II programs by the presence of *HLA-DRA* and *CD74*. We then calculated the mean usage of each gene program for each patient and compared these usage values across patient groups.

To quantify the EM score in bulk transcriptional datasets, we obtained gene expression matrices from each individual study (**Supplementary Table 5**). EM scores were calculated as the sum of the top 10 shared MS1 and NS1 genes (*S100A8, S100A9, S100A12, PLBD1, SELL, HP, GCA, ALOX5AP, MCEMP1,* and *TSPO*) subtracted by the sum of the top 10 MHC-II genes (*HLA-DRA, CD74, HLA-DPB1, HLA-DQA1, HLA-DRB1, HLA-DPA1, HLA-DQB1, EEF1A1, HLA-DRB5,* and *HLA-DMA*). Serum protein abundance measurements (Olink Proteomics) and patient level clinical data from the COVACTA clinical trial were also obtained from a prior publication (*32*).

Meta-analysis across datasets were performed as follows: for each contrast, we first computed the standardized median difference (difference in means normalized by pooled standard deviation) for each dataset, and calculated 95% confidence intervals by bootstrapping the data. Summary effect sizes were then calculated by calculating a weighted average (based on the number of patients for each study) of each effect size.

### HSPC differentiation, culture, and flow cytometry

For each experiment, CD34+ cells were first thawed and rested for 48 hours in SFEM II with 75 nM StemRegenin 1, 3.5 nM UM729, 1X CC110 (StemCell Technologies), and 1X penicillin-streptomycin (Gibco). To initiate differentiation, cells were cultured in the same medium supplemented with 20% pooled human serum (SeraCare) with or without cytokines at the indicated concentrations for each experiment. Recombinant human IL-24, OSM, IL-33, RETN, OPG, TRAIL, TNFβ, IL-6, IL-10, GM-CSF, M-CSF, and G-CSF were obtained from Peprotech. To assess myelopoiesis, cells were stained with the following panel: CD14-FITC (clone M5E2), CD15-APC (clone W6D3), CD11b-AF700 (clone ICRF44), CD34-BV650 (clone 561), HLA-DR-PE/Cy7 (clone L243) (BioLegend) and IL1R2-PE (clone 34141; Thermo). After staining, cells were resuspended in FACS buffer with 5% CountBright beads (Invitrogen) to allow determination of absolute counts during analysis. Flow cytometry data were acquired on an LSR Fortessa (BD Biosciences) and analyzed using FlowJo v10.10 and GraphPad Prism 10.

### COVACTA clinical trial

Details of the COVACTA study have been previously published (*32*, *47*). Briefly, 438 hospitalized COVID-19 patients, confirmed with a positive SARS-CoV-2 PCR test, were randomized 2:1 for anti-interleukin-6 receptor antibody, tocilizumab, or placebo. The NIH COVID-19 ordinal severity scale was used to assess disease severity (3 = hospitalized not requiring supplemental O_2_, 4 = supplemental O_2_, 5 = non-invasive/high flow O_2_, 6 = mechanical ventilation, 7 = mechanical ventilation + additional organ support). Serum protein measurements were obtained using the Olink Explore platform (Olink), and RNA-seq libraries were generated using the TruSeq Stranded mRNA Library Prep kit (Illumina).

### scRNA-seq and data analysis

scRNA-seq combined with cell hashing (*48*) was performed as previously described (*15*). Briefly, cells from multiple culture conditions were labeled with hashtag oligo (HTO) antibodies (BioLegend) and loaded on the Chromium platform using the 3′ v3 profiling chemistry (10X Genomics). Libraries were sequenced to a depth of ∼25,000 reads per cell on a NovaSeq S4 (Illumina). The data were aligned to the GRCh38 reference genome using cellranger v3.1 (10X Genomics). Single-cell data analysis was performed using scanpy with the same preprocessing and filtering parameters described in a prior publication (*15*). cNMF was performed using the same parameters detailed above. MS1, NS1, and MHC-II scores were calculated for the top 30 genes in each module (**Supplementary Tables 2-3**) using the “score_genes” function on scanpy (ctrl_size = 50, n_bins = 25). To analyze mouse scRNA-seq experiments, we first converted the gene expression matrix using mousipy v0.1.5.

### Influenza infection in vivo

Wild-type C57BL/6 mice and Ccr2-/- mice were purchased from Jackson Laboratories or Charles River Hollister. Mice were infected intranasally with the indicated dose of influenza H1N1 A/PR/8/1934 in 50 μL Dulbecco’s Modified Eagle Medium (DMEM) with 2 μg/mL TCPK Trypsin (Thermo) under anesthesia via intraperitoneal injection of 75-80 mg/kg Ketamine and 7.5-15 mg/kg Xylazine. At day 7 post-infection, mice were anesthetized with isoflurane and 500 μL blood was collected by retro-orbital bleeding. After blood collection, mice were euthanized by cervical dislocation while under anesthesia. BALF samples were collected post-euthanasia by instillation and withdrawal of 1 mL PBS into the lung. BALF cells were obtained by centrifugation for 5 min at 400 g, and supernatants were collected for cytokine, sRAGE, and albumin measurements. Neutrophil depletion was performed by intraperitoneal injection of 10 mg/kg anti-Gr1 (clone RB6-8C5, Genentech) on day 1 or day 3 post-infection, and every 2 days thereafter. Mice were treated with 2.5 mg/kg anti-GCSF (clone 67604, ThermoFisher) and/or 20 mg/kg anti-IL-6R (Genentech) at day 1 post-infection, and every 2 days thereafter. Studies were performed in compliance with the regulations of the Association of Assessment and Accreditation of Laboratory Animal Care.

### Cytokine and lung injury measurements

Cytokine levels were measured using the Milliplex Mouse Panel 32-plex (Millipore Sigma). Enzyme-linked immunosorbent assay (ELISA) kits were used to assay mouse albumin (Abcam) and mouse sRAGE (R&D).

### Viral load quantification

MDCK-II cells (ECACC) were cultured in EMEM and seeded (1.5-2×10^4^ cells/well) in black clear-bottom 96-well plates (Corning) to create a monolayer on the following day of viral infection. Five-fold serial dilutions of mouse BALF were carried out in high glucose DMEM (Glucose 4500 mg/L, 0.2% BSA, 1M HEPES, 100 U/mL Penicillin, 100 ug/mL Streptomycin, 2 mM Glutamax (ThermoFisher Scientific) with 2 μg/mL TCPK Trypsin (Sigma). Mouse BALF dilutions were then added to the cells followed by incubation at 37°C for 16-18 h. Cells were then fixed with 100% ice-cold ethanol, blocked with 2% BSA in PBS, and stained with a mouse anti-Influenza A nucleoprotein (Millipore MAB8258). The 96-well plates were then imaged with the Image Express Confocal HT.ai High Content Imaging System (Molecular Devices). Six images of each well were taken followed by quantitative analysis with the MetaExpress 6.0 Program. The dilution which yielded an average of 5-30 positive AF488 events (pfu) was selected and the viral titer in pfu/ml was calculated.

## ACKNOWLEDGEMENTS

This study was supported with funding from Roche, Inc. and in part with federal funds from the Department of Health and Human Services; Administration for Strategic Preparedness and Response; Biomedical Advanced Research and Development Authority, under OT number: HHSO100201800036C. The contract and federal funding are not an endorsement of the study results, product, or company. Schematics created with Biorender.com.

## AUTHOR CONTRIBUTIONS

Experiments: K.K., D.Y., G.L.S., H.L-H., J.K., C.C, M.D.S., and M.R.; Computational Analysis: H.S. and M.R.; Data Curation: H.S., R.N.B., C.M.R., and M.R.; Supervision: S.D., R.N.B., C.M.R., S.B.K., M.X., and M.R.; Conceptualization: C.M.R., S.B.K., M.X., and M.R.; Manuscript Writing: K.K., G.L.S., and M.R.; Manuscript Review and Editing: all authors.

## DATA AVAILABILITY

External datasets analyzed and their corresponding accession IDs are listed in **Supplementary Tables 1 and 5**. scRNA-seq datasets generated in this study will be made available upon publication.

## SUPPLEMENTARY INFORMATION

**Supplementary Figure 1.**
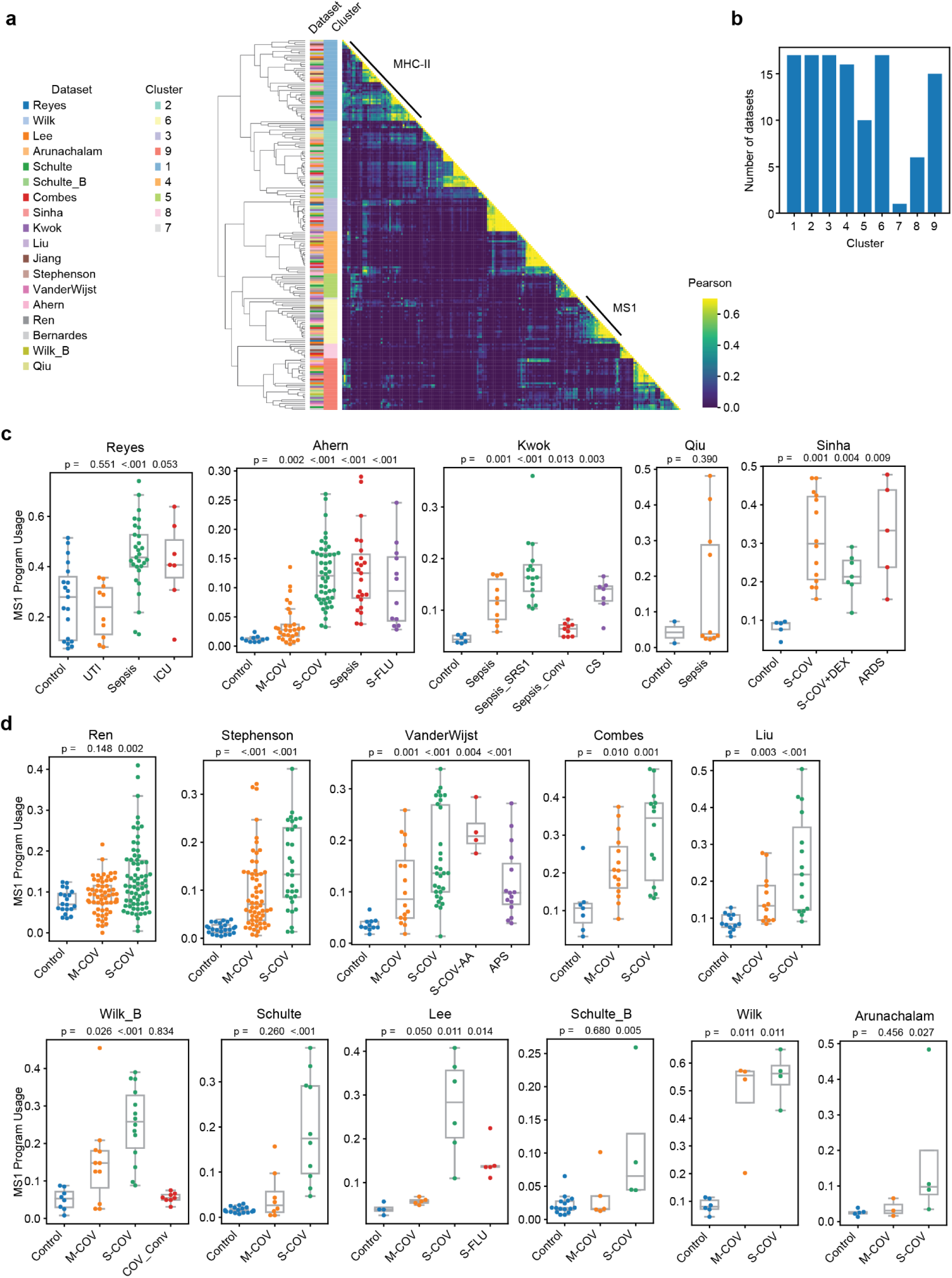
Meta-analysis of monocytes from sepsis and COVID-19 blood scRNA-seq datasets. (a) Correlation matrix of the gene weights (z scores) for the monocyte gene expression programs across 18 scRNA-seq datasets for cohorts of patients with bacterial sepsis or COVID-19. Gene expression modules were derived in an unbiased manner from each dataset using consensus non-negative matrix factorization (cNMF). (b) Barplot showing the number of datasets found in each gene program cluster in (a). (c-d) Mean MS1 program usage in monocytes of bacterial sepsis (c) or COVID-19 (d) patients. Boxes show the median and interquartile range (IQR) for each patient cohort, with whiskers extending to 1.5 IQR in either direction from the top or bottom quartile. p values shown are calculated by comparing each disease state with Control using a two-tailed Wilcoxon rank-sum test. Detailed descriptions of the patient cohorts and numbers of cells and patients for each dataset are described in the corresponding publications (*9*, *10*, *15–28*) and listed in **Supplementary Table 1**. Control, healthy controls; UTI, urinary tract infection with leukocytosis; ICU, intensive care patients without sepsis; Sepsis_SRS1, sepsis patients in the SRS1 subphenotype; Sepsis_Conv, convalescent sepsis; CS: cardiac surgery patients; ARDS, non-COVID-19 acute respiratory distress syndrome; M-COV, mild COVID-19; S-COV, severe COVID-19; S-COV-AA, severe COVID-19 with confirmed type I IFN autoantibodies; S-COV, severe COVID-19; S-COV-AA, severe COVID-19 with confirmed type I IFN autoantibodies; S-COV+DEX, severe COVID-19 treated with dexamethasone; APS, autoimmune polyglandular syndrome; COV_Conv, convalescent COVID-19; S-FLU, severe influenza A.

**Supplementary Figure 2.**
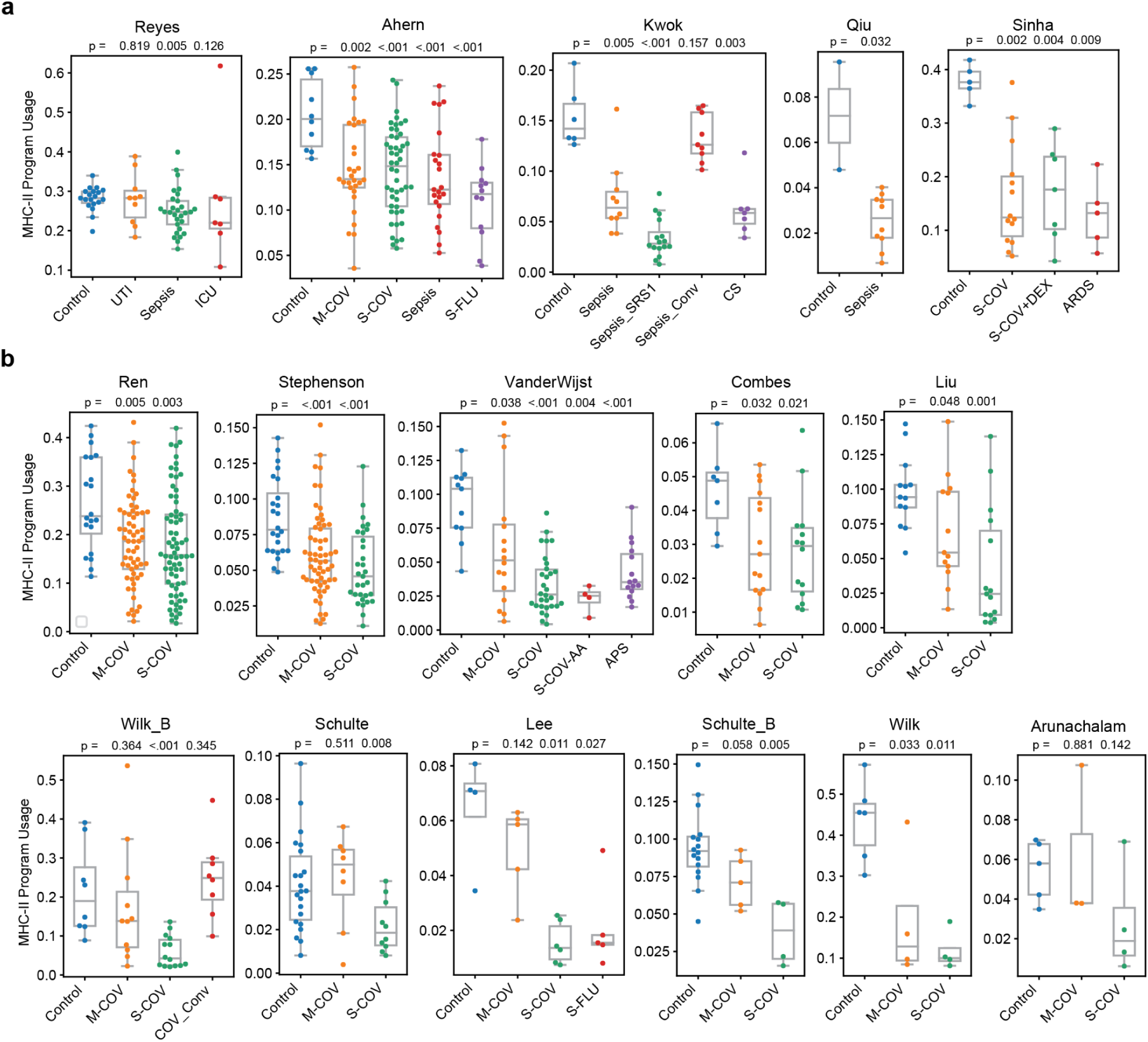
MHC-II program usage in blood monocytes from sepsis and COVID-19 scRNA-seq datasets. (a-b) Mean MS1 program usage in monocytes of bacterial sepsis (a) or COVID-19 (b) patients. Boxes show the median and interquartile range (IQR) for each patient cohort, with whiskers extending to 1.5 IQR in either direction from the top or bottom quartile. p values shown are calculated by comparing each disease state with Control using a two-tailed Wilcoxon rank-sum test. Detailed descriptions of the patient cohorts and numbers of cells and patients for each dataset are described in the corresponding publications(*9*, *10*, *15–28*) and listed in **Supplementary Table 1**. Control, healthy controls; UTI, urinary tract infection with leukocytosis; ICU, intensive care patients without sepsis; Sepsis_SRS1, sepsis patients in the SRS1 subphenotype; Sepsis_Conv, convalescent sepsis; CS: cardiac surgery patients; ARDS, non-COVID-19 acute respiratory distress syndrome; M-COV, mild COVID-19; S-COV, severe COVID-19; S-COV-AA, severe COVID-19 with confirmed type I IFN autoantibodies; S-COV, severe COVID-19; S-COV-AA, severe COVID-19 with confirmed type I IFN autoantibodies; S-COV+DEX, severe COVID-19 treated with dexamethasone; APS, autoimmune polyglandular syndrome; COV_Conv, convalescent COVID-19; S-FLU, severe influenza A.

**Supplementary Figure 3.**
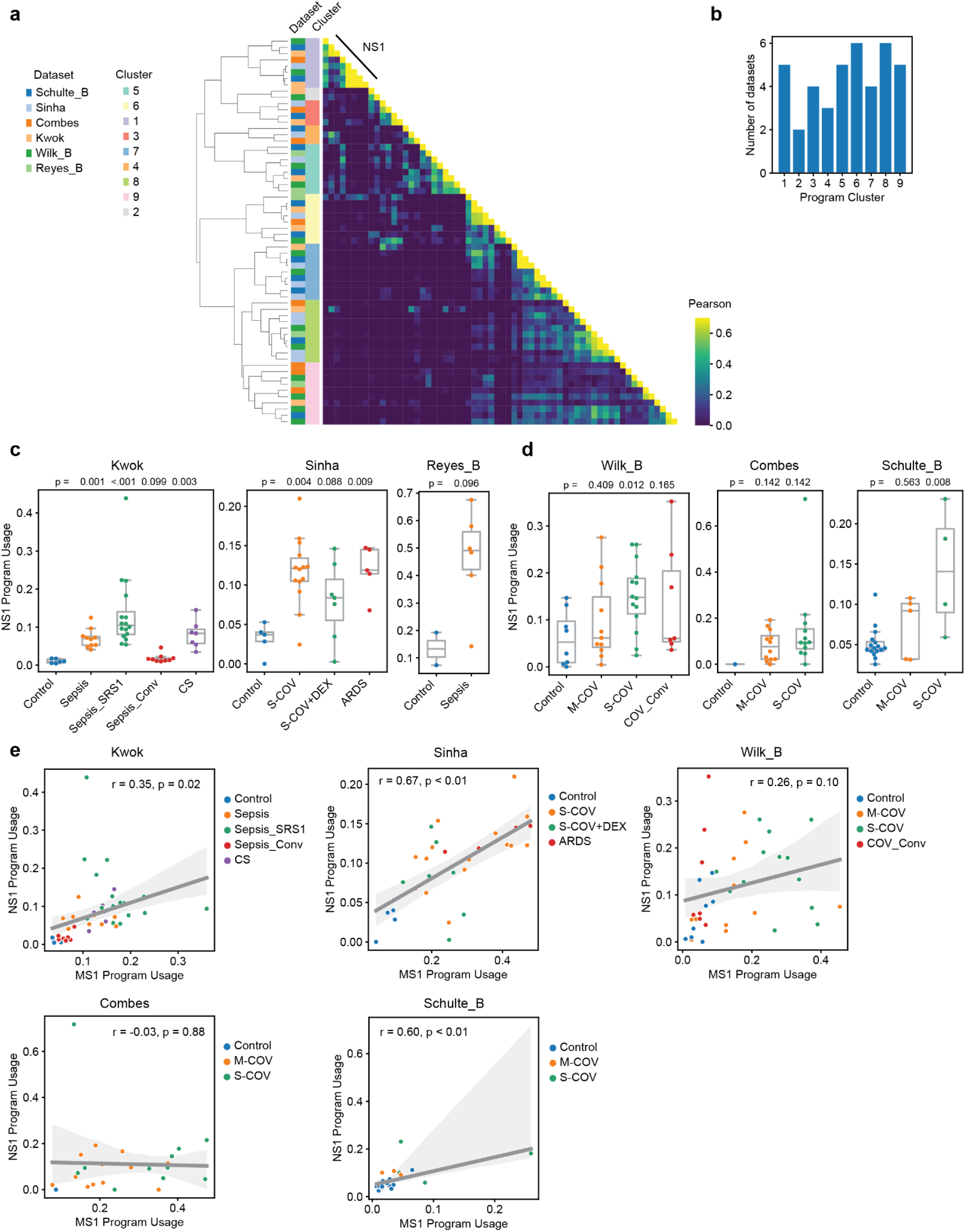
Meta-analysis of neutrophils from sepsis and COVID-19 blood scRNA-seq datasets. (a) Correlation matrix of the gene weights (z scores) for the neutrophil gene expression programs across 18 scRNA-seq datasets for cohorts of patients with bacterial sepsis or COVID-19. Gene expression modules were derived in an unbiased manner from each dataset using consensus non-negative matrix factorization (cNMF). (b) Barplot showing the number of datasets found in each gene program cluster in (a). (c-d) Mean NS1 program usage in neutrophils of bacterial sepsis (c) or COVID-19 (d) patients. Boxes show the median and interquartile range (IQR) for each patient cohort, with whiskers extending to 1.5 IQR in either direction from the top or bottom quartile. p values shown are calculated by comparing each disease state with Control using a two-tailed Wilcoxon rank-sum test. (e) Scatterplot showing correlation between mean monocyte MS1 and neutrophil NS1 module usage for each patient. Line and shadow indicate linear regression fit and 95% confidence interval, respectively. Significance of the correlations (Pearson r) were calculated with a two-sided permutation test and corrected for multiple comparison of modules. Detailed descriptions of the patient cohorts and numbers of cells and patients for each dataset are described in the corresponding publications (*9*, *15*, *17–19*, *24*) and listed in **Supplementary Table 1**. Control, healthy controls; UTI, urinary tract infection with leukocytosis; ICU, intensive care patients without sepsis; Sepsis_SRS1, sepsis patients in the SRS1 subphenotype; Sepsis_Conv, convalescent sepsis; CS: cardiac surgery patients; ARDS, non-COVID-19 acute respiratory distress syndrome; M-COV, mild COVID-19; S-COV, severe COVID-19; S-COV, severe COVID-19; S-COV+DEX, severe COVID-19 treated with dexamethasone; COV_Conv, convalescent COVID-19.

**Supplementary Figure 4.**
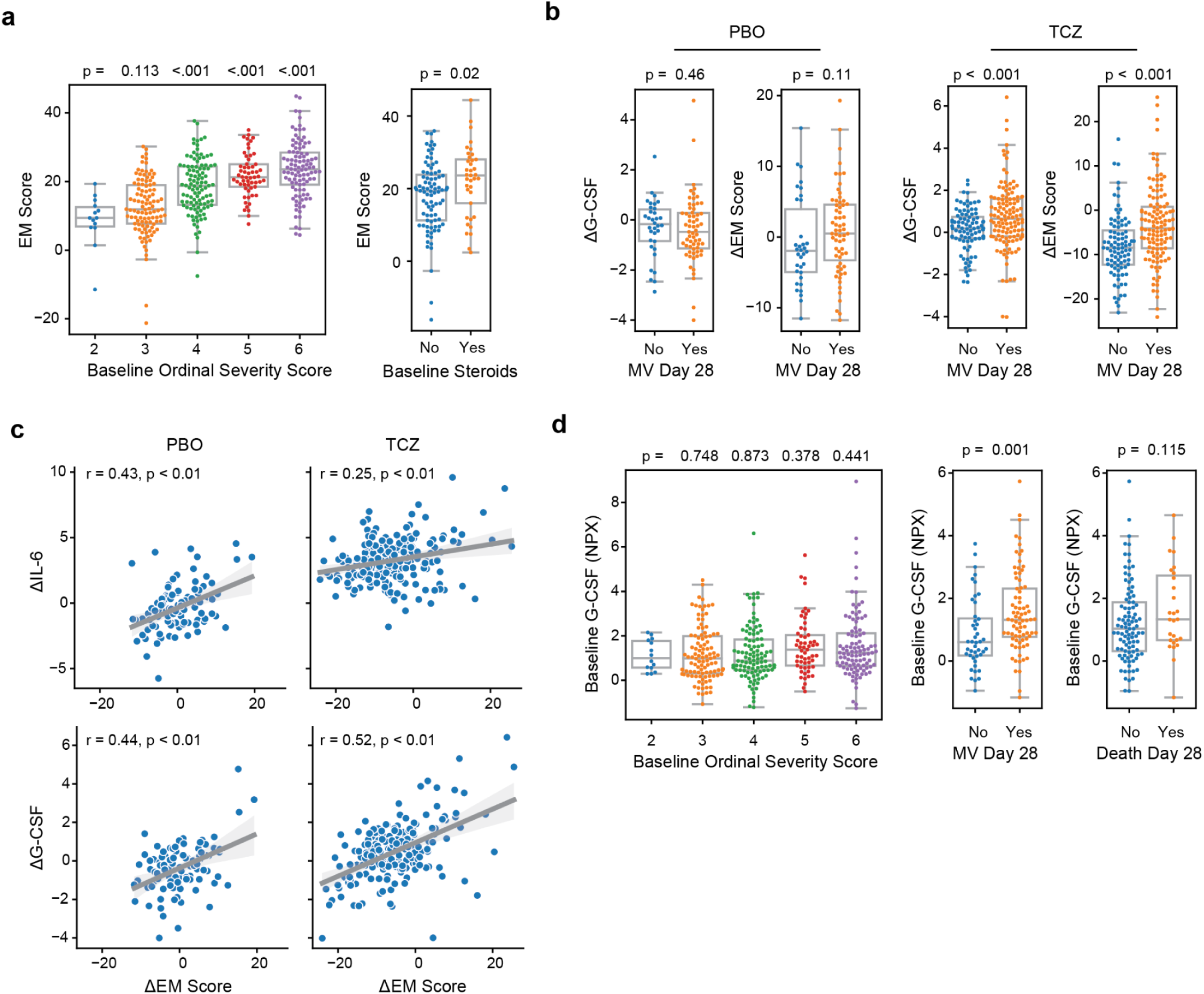
Emergency myelopoiesis scores and G-CSF levels in the COVACTA clinical trial. (a) EM scores in patients from COVACTA grouped by baseline ordinal score (left) or steroid use (right). p values shown are calculated for each comparison using a two-tailed Wilcoxon rank-sum test. (b) Changes (baseline to day 3) in EM score or G-CSF levels in patients from the PBO (left) or TCZ (right) arm of COVACTA grouped by mechanical ventilation status on day 28. p values shown are calculated for each comparison using a two-tailed Wilcoxon rank-sum test. (c) Scatterplots showing correlations between changes (baseline to day 3) in IL-6 (top) or G-CSF (bottom) and EM scores in PBO (left) or TCZ (right) arms. Line and shadow indicate linear regression fit and 95% confidence interval, respectively. Significance of the correlations (Pearson r) were calculated with a two-sided permutation test and corrected for multiple comparison of modules. (d) G-CSF in patients from COVACTA grouped by baseline ordinal score (left) or steroid use (right). p values shown are calculated for each comparison using a two-tailed Wilcoxon rank-sum test. (d) Baseline G-CSF levels in patients from COVACTA grouped by baseline ordinal score (left), day 28 mortality (middle) or day 28 mechanical ventilation status (right). p values shown are calculated for each comparison using a two-tailed Wilcoxon rank-sum test. Boxes show the median and interquartile range (IQR) for each patient cohort, with whiskers extending to 1.5 IQR in either direction from the top or bottom quartile. NPX, normalized protein expression.

**Supplementary Figure 5.**
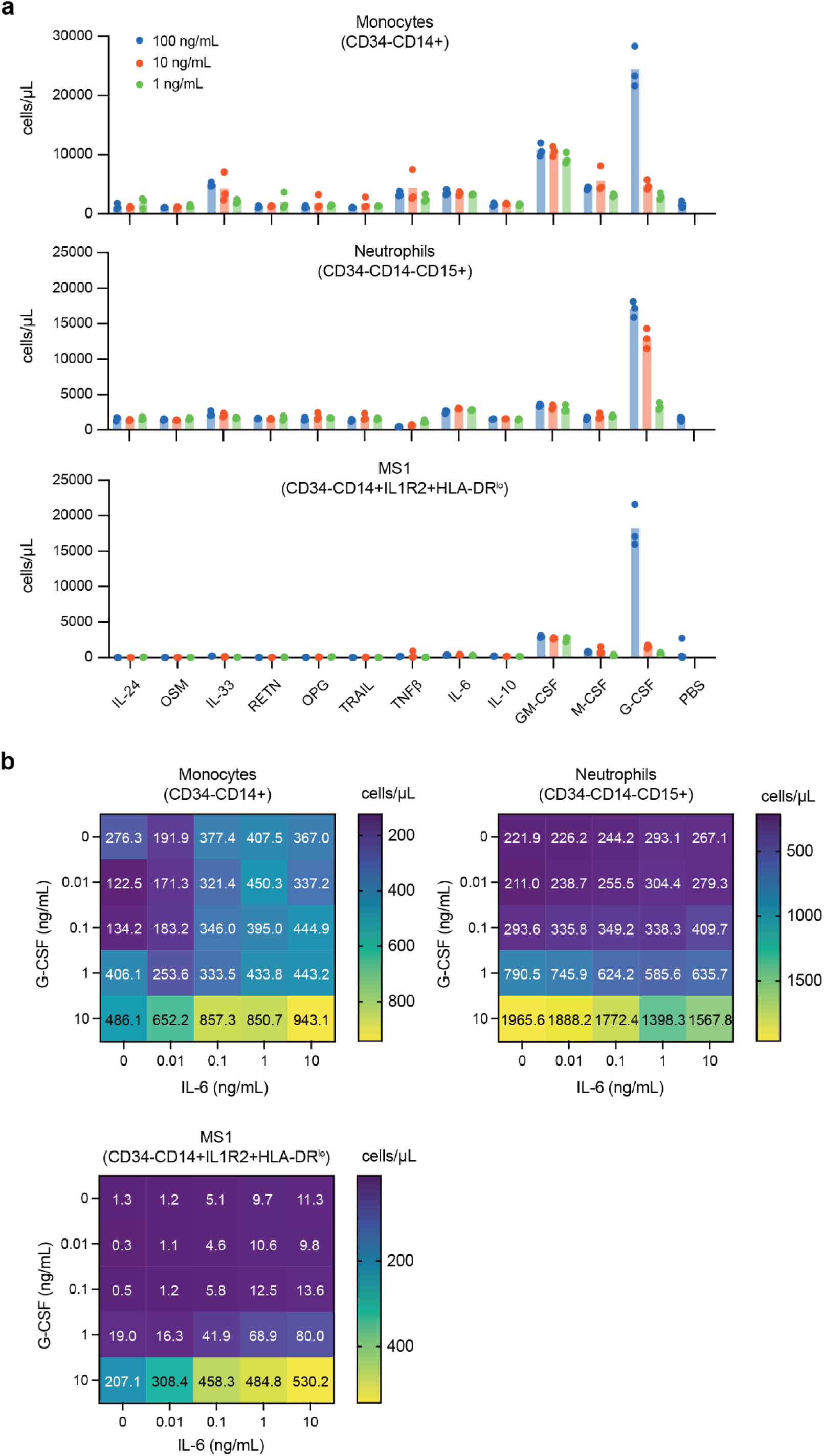
Effects of cytokine on MS1 monocyte and neutrophil production *in vitro*. (a) Number of monocytes (top), neutrophils (middle), and MS1 monocytes (bottom) generated from HSPCs incubated with each cytokine at the indicated concentration. (b) Number of monocytes (top), neutrophils (middle), and MS1 monocytes (bottom) generated from HSPCs incubated with different concentrations of IL-6 (x-axis) and G-CSF (y-axis).

**Supplementary Figure 6.**
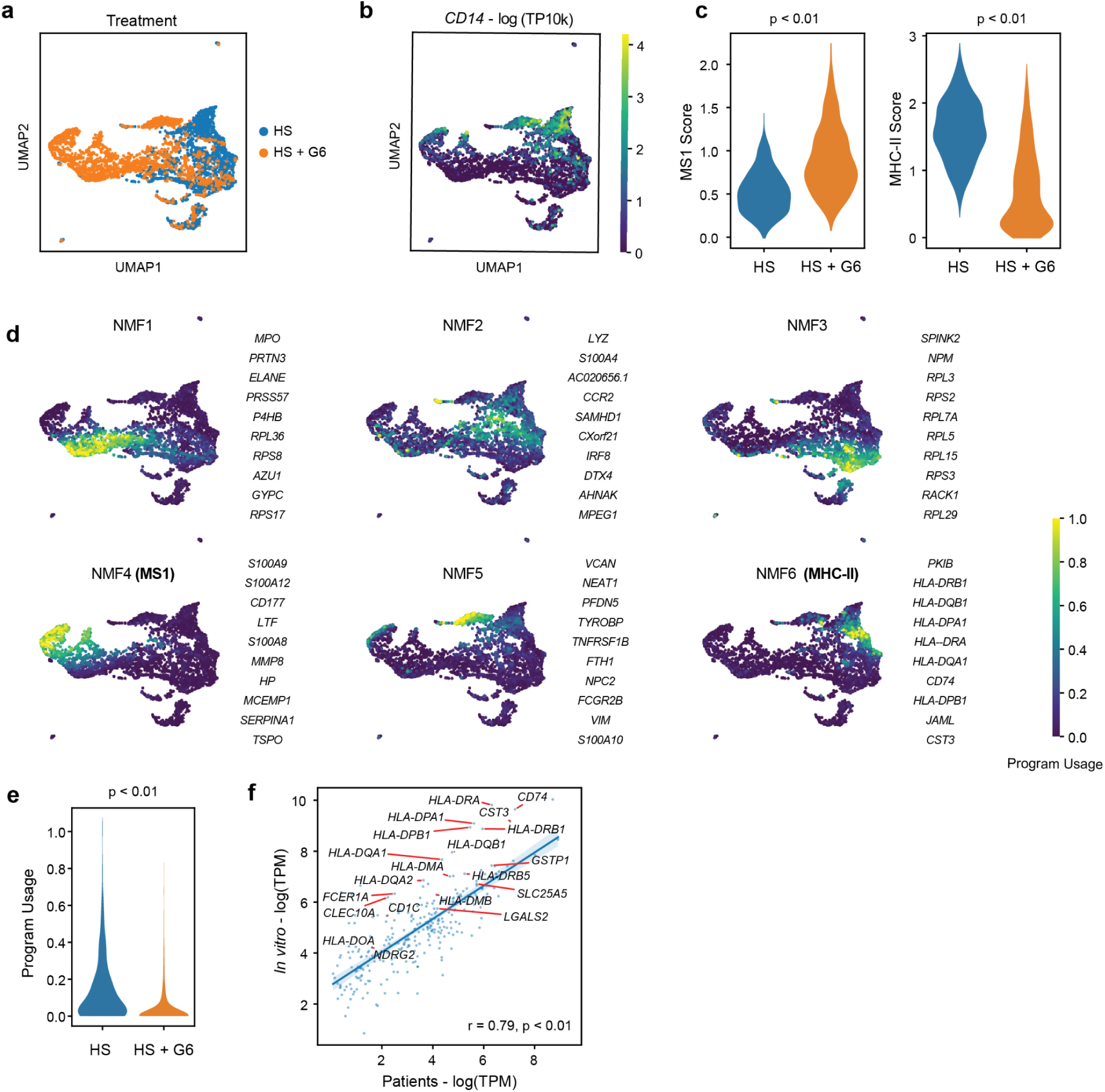
scRNA-seq of HSPCs differentiated with IL-6 and G-CSF. (a-b) UMAP of scRNA-seq data from HSPCs treated with healthy serum (HS) or healthy serum with 10 ng/mL IL-6 and G-CSF (HS+G6). Colors indicate treatment (a) or CD14 expression (b) for each cell. (c) Violin plots showing MS1 (left) and MHC-II (right) scores in CD14+ monocytic cells across treatments. p values shown are calculated for each comparison using a two-tailed Wilcoxon rank-sum test. (d) UMAP of scRNA-seq data from the experiment in (a). Colors indicate usage of each gene program. The top 10 genes in each program are indicated on the right. (e) Violin plot of MHC-II gene program usage in monocytes differentiated from HSPCs using the indicated condition. (f) Gene weight correlation between the MHC-II program detected in patients (x-axis) and the MHC-II program detected in differentiated HSPCs (y-axis). Significance of the Pearson correlations (r) was calculated with a permutation test. The top 20 genes with the highest z-score loadings in the patient MHC-II program are labeled. UMAP, uniform manifold approximation projection.

**Supplementary Figure 7.**
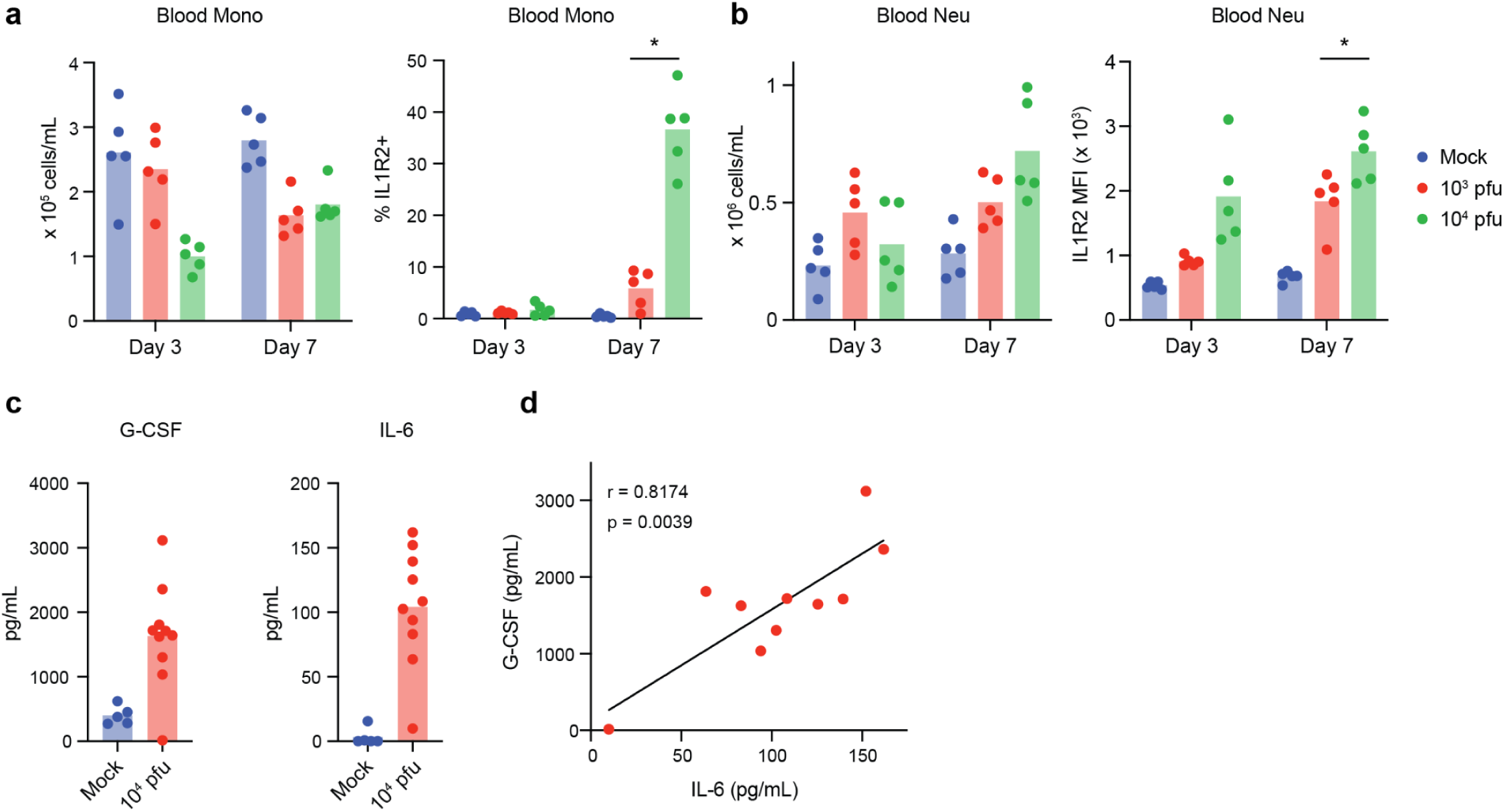
Emergency myelopoiesis during severe influenza infection *in vivo*. (a) Number (left) and fraction of IL1R2+ (right) BALF monocytes in mice infected with PR8 at the indicated time point. (b) Number (left) and IL1R2 mean fluorescence intensity (MFI; right) of BALF neutrophils in mice infected with PR8 at the indicated timepoint. (c) Blood G-CSF (left) and IL-6 (right) levels in mice infected with 10^4^ pfu PR8 at day 7 post-infection. (d) Correlation between blood G-CSF (y-axis) and IL-6 (x-axis) levels in mice infected with 10^4^ pfu PR8 at day 7 post-infection. Significance of the Pearson correlation (r) was calculated with a permutation test.

**Supplementary Figure 8.**
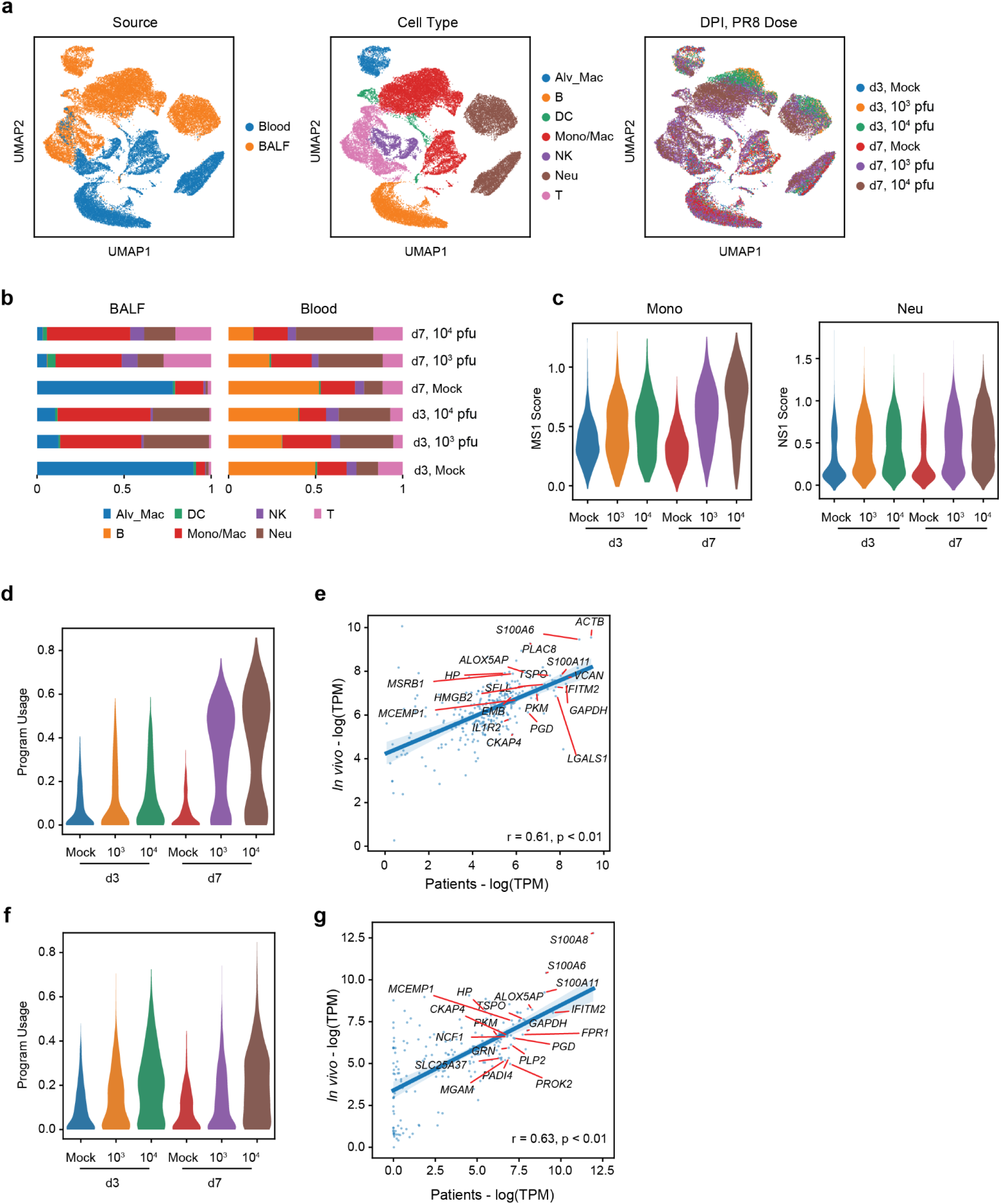
scRNA-seq of blood and BALF immune cells from influenza-infected mice. (a) UMAP of scRNA-seq data from blood and BALF immune cells of mice infected with PR8. Colors indicate tissue source (left), cell type (middle), and dose/timepoint (right) for each cell. (b) Fractional abundance of each cell type in the BALF (left) and blood (right) for the indicated dose/timepoint. (c) Violin plots showing blood monocyte MS1 (left) and neutrophil NS1 (right) scores for the indicated dose/timepoint. (d,f) Violin plots of mouse MS1 (d) and NS1 (f) program usage in mice infected with PR8. (e,g) Gene weight correlation between the MS1 (e) and NS1 (g) gene programs detected in patients (x-axis) and in mice (y-axis). Significance of the Pearson correlations (r) was calculated with a permutation test. The top 20 genes with the highest z-score loadings in each patient program are labeled.

**Supplementary Table 1.** List of scRNA-seq datasets analyzed

**Supplementary Table 2.** Top genes associated with monocyte consensus programs

**Supplementary Table 3.** Top genes associated with neutrophil consensus programs

**Supplementary Table 4.** Effect sizes of EM gene programs across scRNA-seq datasets

**Supplementary Table 5.** List of bulk RNA-seq datasets analyzed

**Supplementary Table 6.** Effect sizes of EM scores across bulk RNA-seq datasets

**Supplementary Table 7.** Correlations between changes in protein levels and EM scores

